# Addressing multiple sources of uncertainty in the estimation of parrot abundance from roost counts: a case study with the Vinaceous-breasted Parrot (*Amazona vinacea*)

**DOI:** 10.1101/455774

**Authors:** Viviane Zulian, Eliara Solange Müller, Kristina L. Cockle, Arne Lesterhuis, Roberto Tomasi Júnior, Nêmora Pauletti Prestes, Jaime Martinez, Gonçalo Ferraz

## Abstract

Population size is a key predictor of extinction risk and is critical to listing species in IUCN threat categories. The population size of parrots—one of the most threatened bird families—is often assessed using roost counts, which suffer from multiple sources of uncertainty that need to be addressed in monitoring efforts. To improve estimates of abundance for endangered Vinaceous-breasted Parrot (*Amazona vinacea*), we compared extensive roost counts over the whole range of the species (Argentina, Paraguay, Brazil) with an intensive regional survey designed to address five sources of uncertainty about parrot abundance in western Santa Catarina state (WSC), Brazil, in 2016 and 2017. We estimated regional-scale abundance using a sampling design that minimizes double counting and an N-mixture model of replicated count data, which accounts for imperfect detection, implemented in a Bayesian framework. The whole-range counts amounted to 3,888 and 4,084 individuals in 2016 and 2017, respectively; regional estimates were 945 ± 50 and 1,393 ± 40 individuals, for the same two years. We found no evidence of population growth because the increase in numbers matched an increase in observation effort on both spatial scales. When extrapolating the WSC abundance estimate to three hypothetical geographical range areas of the species, under the simplifying assumption of homogenous density, we obtained values above the whole-range counts, but within the same order of magnitude, putting the global population size of Vinaceous-breasted Parrot in the thousands of individuals. Although our estimates of abundance and geographic range are larger than those currently reported by the IUCN, we suggest that Vinaceous-breasted Parrot remain in the ‘Endangered’ IUCN threat category pending further investigation of population trends. We recommend that roost-monitoring programs for parrots consider and address sources of uncertainty through field protocols and statistical analysis, to better inform assessments of population size, trends, and threat status.

---

Population size is arguably the most important state variable in population biology (Gaston 1994); along with range size, it is the most evident predictor of extinction risk (Lawton 1995) and plays a central role in the assessment of any population management strategy (Caughley 1994, Norris 2004). Abundance is also directly implicated in three of the five IUCN (International Union for the Conservation of Nature) criteria for listing species in threat categories (Mace *et al.* 2008). Among the animal groups in most urgent need of abundance information, parrots (Psittaciformes) stand out for having the highest number of threatened species of all non-passerine bird orders (Olah *et al.* 2016). Of the 398 extant species of parrots, 112 (28%) are listed as threatened (i.e. Vulnerable, Endangered, or Critically Endangered); of these, 88 are listed as declining by the IUCN (BirdLife International 2017). The key causes of parrot population decline are habitat loss (due to deforestation and agroindustrial expansion) and nest poaching (due to the illegal pet trade; Wright *et al.* 2001, Olah *et al.* 2016, Berkunsky *et al.* 2017).

While threatened species do not recognize national borders, environmental regulations do, so it is reasonable to ask which countries can contribute most to protecting any given group. Brazil has the highest number of globally threatened parrot species (Olah *et al.* 2016), and among these, one of the least known, both in terms of organismal biology and population dynamics, is the Vinaceous-breasted Parrot (*Amazona vinacea*). Currently listed as ‘Endangered’, the Vinaceous-breasted Parrot is restricted to the Atlantic Forest biome, with a geographic range that falls mostly within Brazil, with small areas of occurrence in the Argentinian province of Misiones and in eastern Paraguay (Cockle *et al.* 2007, Carrara *et al.* 2008, Segovia and Cockle 2012, Prestes *et al.* 2014; Fig. 1). Vinaceous-breasted Parrots appear to be associated with the ancient Paraná Pine (*Araucaria angustifolia*; Collar *et al.* 2017, Tella et al. 2016), but they are also frequently observed foraging and nesting in other trees (Cockle *et al.* 2007, Prestes *et al.* 2014, Bonaparte and Cockle 2017); furthermore, their rather uncertain geographic range seems to extend well beyond the current range of Paraná Pine forests (Cockle *et al.* 2007, Carrara *et al.* 2008). Uncertainty about the geographic range of the Vinaceous-breasted Parrot results in large part from parrot movements, which appear motivated by temporal variation in food availability. Seasonal movements are reported in coincidence with fruiting of *Ocotea puberula*, *Podocarpus lambertii*, *Vitex megapotamica*, Juçara palms (*Euterpe edulis*), and Paraná Pine (Collar et al. 1992, Forshaw 2010, Prestes et al. 2014). Unpredictable movements make it difficult to anticipate where the birds are, or whether parrots seen in different places are the same or different individuals, presenting interesting challenges to the estimation of population size.

**Figure 1.**
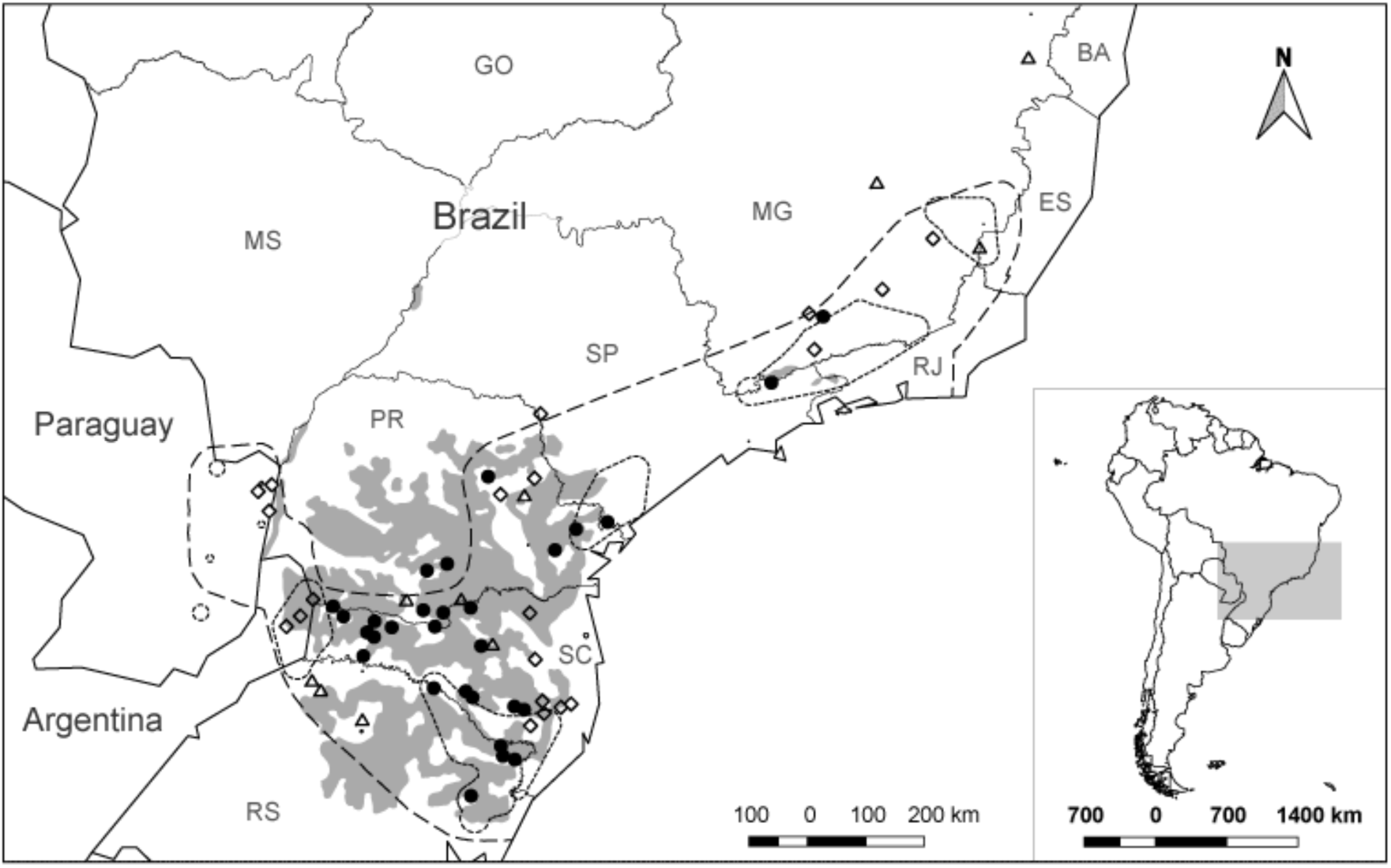
Count sites and hypothetical Vinaceous-breasted Parrot range areas used for extrapolated estimates of population size. Symbols represent sites visited only in 2016 (diamonds), only in 2017 (triangles), or in both years (filled circles). Outlines represent the IUCN ‘Extant’ range (dotted line), the IUCN ‘Possibly Extant’ range (dashed line) and the potential extent of *Araucaria angustifolia* forests (gray polygon). For ease of representation, sites located less than fifteen kilometers from each other are shown by only one symbol. Uppercase letter pairs stand for Brazilian states of Mato Grosso do Sul (MS), Minas Gerais (MG), Espírito Santo (ES), São Paulo (SP), Rio de Janeiro (RJ), Paraná (PR), Santa Catarina (SC) and Rio Grande do Sul (RS).

According to information compiled by the IUCN, the extant geographic range of the Vinaceous-breasted Parrot covers an area of approximately 145,700 square kilometers (BirdLife International and Handbook of the Birds of the World 2016; Fig. 1). This range consists of five patches with tens of thousands of square kilometers each, and eleven patches of a few hundred square kilometers. The discontinuity reflects not only the species’ true range, but also the scarcity of information about population structure and movements. Indeed, the IUCN recently updated the range map with a larger, ‘possibly extant’ layer that encloses all of the patches above (Fig. 1).

Part of the challenge in understanding the distribution and abundance of the Vinaceous-breasted Parrot comes from its life cycle. Our field observations prior to this work suggest that breeding individuals disperse in pairs throughout the species’ range between July and December (VZ, unpublished data). Towards the end of the breeding season, from December to January, they start congregating every evening in roosts that they may or may not use throughout the entire non-breeding period. The number of roosting individuals can vary from fewer than ten to hundreds of individuals both among roosts and among days at the same roost during the January-June non-breeding period (VZ, unpublished data). When August begins, there are virtually no parrots left at the roosts and the population is once again dispersed across hundreds of nesting sites. Despite the difficulties inherent to locating roosts and counting the number of individuals at roosts, roost counts are at present the most effective way of assessing the population size and delimiting the range of the species.

Roost counts can be designed in many different ways but they always involve locating roosts, choosing the appropriate time for counting, and actually counting a number that is as close as possible to the real number of animals present in the area (Casagrande and Beissinger 1997). In order to improve knowledge of the distribution and abundance of the Vinaceous-breasted Parrot from roost counts, one should approach the three tasks of locating, timing, and counting roosts in a way that minimizes five key sources of uncertainty about the end result. The first and second sources have to do with locating roosts. First, there is uncertainty about the extent of the Vinaceous-breasted Parrot’s distribution. When does a gap in the range map represent true absence of the species vs. absence of observations? This problem is well represented by the difference between the IUCN ‘Extant’ and ‘Possibly Extant’ ranges in Figure 1. The second source is uncertainty about density of roosts at the local to regional scale. At what point should one stop spending resources on finding more roosts, versus dedicating time to studying the known roosts in detail? The third source of uncertainty pertains to the movement of individuals between roosts and constrains the timing of counts: if roosts or counting sites correspond to isolated local populations, different roosts could be counted at any time throughout a non-breeding season. If, on the contrary, individuals move between roosts, then such movements have to be accounted for, or counts have to be simultaneous. The fourth and fifth sources of uncertainty relate to the counting technique itself, and address, respectively, false negative and false positive observations of individuals. A false negative happens when a parrot that is present at a site is not counted because it was not seen. A false positive happens when a parrot is counted twice by mistake.

This paper offers an assessment of Vinaceous-breasted Parrot abundance for the years 2016 and 2017. We follow a two-pronged approach that combines data from two spatial scales, two counting techniques, and two research teams. At the whole-range scale, we provide a global count of parrots observed in all Vinaceous-breasted Parrot roosts known to us, throughout the entire range of the species. Whole-range counts offer a lower bound for the population size of the species. At the regional scale, we offer a statistical estimate of the number of Vinaceous-breasted Parrots present in Western Santa Catarina (WSC; Fig. 2) that seeks to address all five sources of uncertainty listed in the previous paragraph. We chose to focus the regional research on WSC because a) being an area of intense agro-industrial activity with no previously published Vinaceous-breasted Parrot observations in the scientific literature, it has been left out of the species’ IUCN Extant map; b) it sits between two important Vinaceous-breasted Parrot habitat areas in different countries (Misiones in Argentina and the Araucaria forests of Eastern Santa Catarina in Brazil), and c) based on our previous experience, we expected to find roost-sites that were not yet documented in WSC. To connect the regional and the whole-range approaches we extrapolate our estimate of abundance in WSC to three different global estimates of population size under the simplifying assumption of homogeneous density and compare the extrapolated result with the whole-range counts. We discuss the implications of our results for listing Vinaceous-breasted Parrot as an endangered species.

## Methods

### Study area

Whole-range sampling took place over 59 sites spanning an area from northern Minas Gerais in the north to northeastern Rio Grande do Sul, in the south. This area extends west to (and includes) eastern Paraguay and the province of Misiones in northeastern Argentina (Figure 1). One quarter (15) of the count sites were located inside the Vinaceous-breasted Parrot IUCN Extant range and the remainder (44) were outside. Sites correspond to regularly used roosts and to points of frequent flyover by parrots at dawn and dusk (Supplemental Material Table S1). Our research team and collaborators identified the count sites, sometimes over decades of Vinaceous-breasted Parrot observation (e.g. Cockle *et al.* 2007, Segovia and Cockle 2012). All sites are located within the Atlantic Forest, defined by the southeast, Atlantic portion of the ‘tropical and subtropical moist broadleaf forest’ eco-region in South America (Olson *et al.* 2001).

The regional-scale study area was the western part of the Brazilian state of Santa Catarina (WSC; Figure 2), a rectangle-shaped area of 34,000 km^2^ (IBGE 2015) extending West-East between the Uruguay river (to the South) and the ridgeline that separates the Uruguay and Iguaçú watersheds (to the North). Although mostly deforested, the area is adjacent to two large patches of forest habitat: the Atlantic Forest of the Argentine Province of Misiones, to the west, and the Araucaria forests of Eastern Santa Catarina, to the east (Figure 2). WSC is remarkable for having a surprisingly high frequency of Vinaceous-breasted Parrot sightings by citizen scientists (Wikiaves 2008) in an area that is almost entirely (88%) outside the IUCN-defined extant range of the species (Figure 1). WSC falls within the Araucaria forest and the Interior forest biogeographic sub-regions of the Atlantic Forest, which have lost, respectively, 87 and 93% of their forest cover since the onset of European colonization (Ribeiro *et al.* 2009). Nowadays, the remaining forest patches in WSC (Figure 2) are surrounded by agro-industrial development, consisting mostly of soybean, eucalyptus (*Eucalyptus* sp.), and pine (*Pinus* sp.) plantations (Fearnside 2001, Baptista and Rudel 2006). The nine WSC sampling sites were a subset of the whole-range sites. They comprised all known Vinaceous-breasted Parrot roosts in WSC and they all coincided with Araucaria forest patches >10 m tall. Four of the nine regional sites (*Guatambu*, *Campo Erê*, *Abelardo Luz* and *Água Doce*) had very open to non-existent vegetation under the Araucaria canopy (Figure 2).

**Figure 2.**
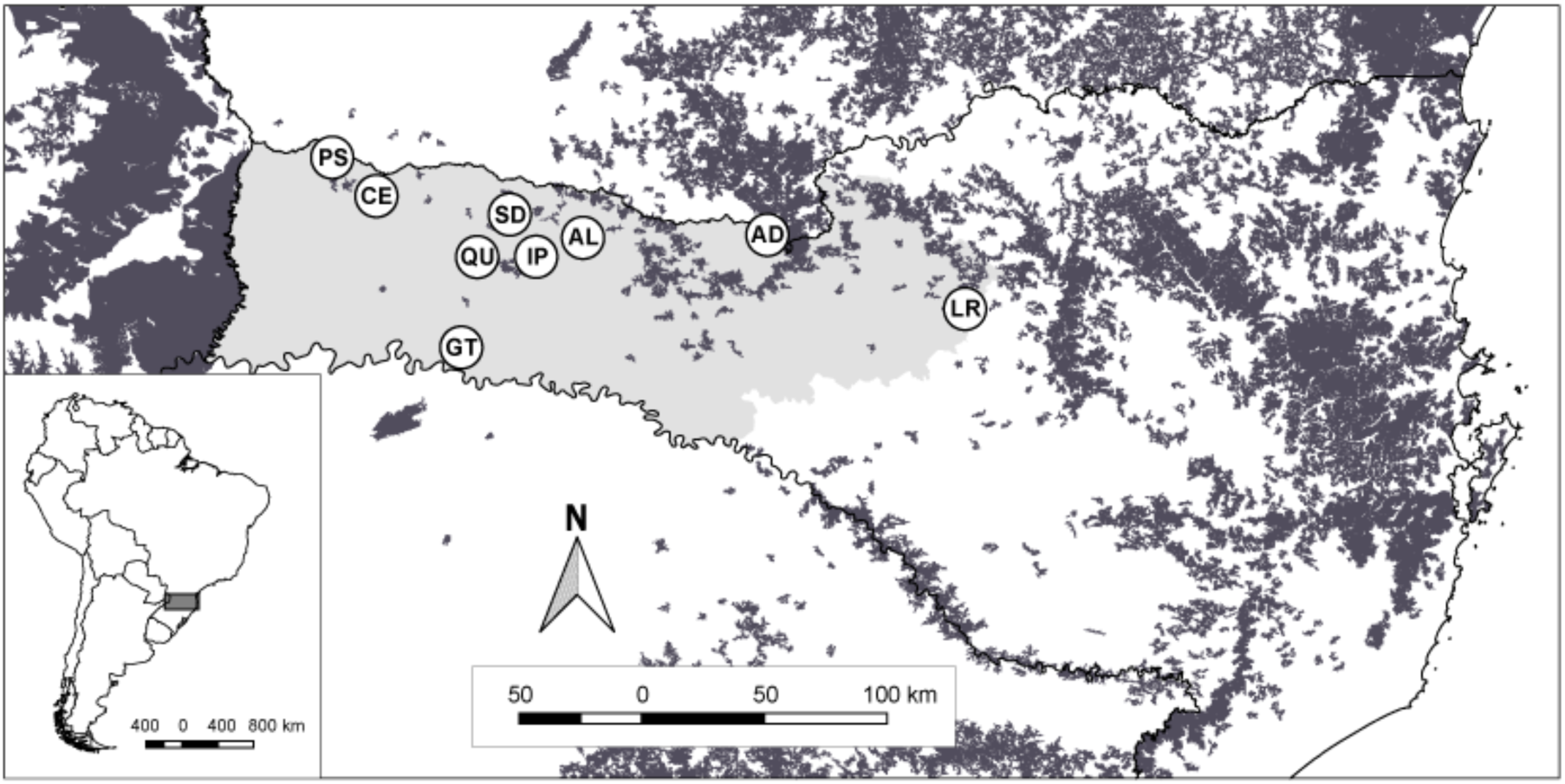
Regional-scale study area of Western Santa Catarina (light gray) and regional forest cover (dark gray). Dark gray areas represent every patch of forest (excluding tree farms) with more than five square kilometers, according to the Brazilian Ministry of the Environment’s *Mapa de Cobertura Vegetal dos Biomas Brasileiros* (MMA 2007). The circles show the location of all presently known WSC roosts with their name abbreviations: PS (*Palma Sola*), CE (*Campo Erê*), GT (*Guatambu*), QU (*Quilombo*), SD (*São Domingos*), IP (I*puaçu*), AL (*Abelardo Luz*), AG (*Água Doce*) and LR (*Lebon Régis*).

### Data collection

Sampling at the whole-range scale was carried out by 26 volunteer teams (Supplemental Material Table S1) coordinated by JM and NPP. Whole-range counts took place on 24-26 March 2016 in Argentina, 29 April to 15 May 2016 in Paraguay and Brazil, and 24 April to 15 May 2017 in Paraguay and Brazil. Each team worked in areas that were familiar to its members, enabling us to cover most of the range in a relatively short period and thus minimize the possibility of double-counting between sampling sites. By minimizing double-counting we reduce the risk of false-positive errors in the assessment of abundance. Of the total 59 sites, 20 were sampled only in 2016, 10 only in 2017, and 29 in both years (Supplemental Material Table S1). We visited sites once per year, counting parrots at the beginning or at the end of the day.

Counts started at dawn (30 minutes before sunrise) or dusk (90 minutes before sunset) and lasted until we could not detect parrot movement into or out of the roost for 20 minutes—which always happened within two hours of the beginning of the count. The number of counting posts at each site varied between one and five, located at strategic points for observing movement of flying parrots in and out of the site area. Each count was performed by a team of one to ten observers who registered the number of parrots arriving or leaving the area, the flight direction, and the time. Whenever there was more than one post in a count, observers from different posts met at the end of the count to compare notes and agree on the minimum number of individual parrots seen.

Fieldwork at the regional-scale was carried out by a single team coordinated by VZ and ESM. Here, we performed monthly visits to count parrots at each site, across two consecutive non-breeding seasons: from December 2015 to July 2016, and from February to June 2017. Having just one team repeating procedures on the same sites allowed for a much tighter control and coordination of field technique at the regional than at the whole-range scale. To minimize the uncertainty about the Vinaceous-breasted Parrot distribution and roost density over the regional-scale, we spent one day per month contacting WSC residents and searching for roosts. The number of sites sampled per month increased from four to nine, as we discovered new roosts throughout the study period (Figure 2). We could identify roosting trees on every site, even though occasionally, on some sites, we only saw parrots flying over, without being able to tell exactly where they perched for the night (e.g. *São Domingos*, *Palma Sola*). Monthly visits lasted from four to eight days, during which we counted the number of individuals present at each site (or roost) between one and four times, with an average of 2.2 counts per roost per visit. To avoid the possibility of a parrot being counted twice in different roosts during the same month, each visit was performed in the shortest period possible. Regional-scale counts were thus obtained in 13 field visits to WSC, eight during the first and five during the second year. We started out by sampling four roosts in December 2015, increased to five in February 2016, seven in May 2016, and nine in May 2017 (Supplemental Material Table S2). The *Lebon Régis* site was only visited during the whole-range count of both years. In total, we completed 179 roost counts at the regional scale.

Regional-scale counts started at dawn or dusk, and lasted until we could not detect parrot movements, following the same times and criteria as prescribed for the whole-range counts. We visited every roost before the first count to establish three observation posts per roost, in strategic locations for observing the arrival and departure of parrots. Each count was performed by a team of three observers (one at each post), each equipped with a roost area map, a compass, an audio recorder, and a radio to communicate with team members about parrots going their way. Every time an observer saw one or more Vinaceous-breasted Parrots, she recorded the number of individuals, the time, and the direction of flight, as well as any other comments that could help understand the movement of the birds. At the end of each count, the team of three observers met to compare their notes and agree on one ‘most reasonable’ (MR) and one ‘highly conservative’ (HC) count result. The difference between MR and HC counts lies in how observers treat the possibility of double counting. Suppose, for example, that an observer sees five parrots arriving at a roost and a few minutes later sees another arrival of three individuals. Based on this information, the MR count is of eight individuals. Suppose further, however, that one of the observers in the trio determined that there were unseen, but heard, parrots leaving the roost during the time between the two observations above. In this case, the team might judge that there was some, however small, possibility that the second group of three was a subset of the first group of five who had left undetected and returned within sight. If that were the case, the HC count result should be five and not eight, because five is the absolute minimum number of birds that the team is sure to have seen arriving at the roost.

The consideration of MR and HC count results addresses one source of uncertainty about Vinaceous-breasted Parrot abundance estimates: the possibility that some animals may be counted more than once. There is, however, a second source of uncertainty that deserves attention, which is imperfect detection, i.e. that some animals are not counted even though they are present at the roost. To address imperfect detection, we replicated our counts by working simultaneously with two teams of three observers, at the same roost and time. We placed two observers (one from each team of three) at each of the posts, keeping sufficient distance between observers to preclude overhearing radio communications. Furthermore, we ensured that observers from different teams did not exchange any information about their observations until each team had separately agreed on its count results. We thus treat every team-specific count of a given roost and month, whether at dawn or dusk, as an independent sample of that roost for that month. The maximum of four counts per roost per month derives from the work of two teams at two different times.

### Data analysis

We statistically modeled the regional-scale data to estimate abundances for each roost site and month using an N-mixture model (Royle 2004). We analyzed MR and HC counts as separate data sets, each summarized by a year-specific array *C* with dimensions *S* by *R* by *M*, where *S* = 9 is the number of roost sites, *R* = 4 is the maximum number of replicate counts per roost in any month, and *M* = 14 is the number of sampling months. Elements *C_ijk_* of this three-dimensional array give the number of parrots counted at the *j*^th^ count of the *i*^th^ roost in the *k*^th^ month, with *i* = 1, …, *S*, *j* = 1, …, *R*, and *k* = 1, …, *M*. The N-mixture model represents the number *N_ik_* of individuals in roost *i* and month *k* as drawn from a Poisson distribution with parameter *λ_k_*. For simplicity, we drop the subscript *k* from the notation below, but we do model each month separately and therefore have monthly estimates of the Poisson parameter and of the number of parrots at each roost. The most straightforward implementation of Royle’s (2004) model accounts for imperfect detection by modeling the counts *C_ij_* as the result of a binomial sample with *N_i_* independent trials and probability of success *p* (which also takes a different value every month). The Binomial distribution, however, equates the probability of success *p* with the probability *p* of detecting one individual parrot and implies that such probability of detection is independent among parrots. This would be reasonable if parrots moved about independently of each other, but they do not; instead, they form groups of variable sizes where large, noisy groups are easier to detect than small groups. To address this problem, we followed Martin *et al.* (2011)’s approach of modeling detection as a Beta-binomial distribution, with parameters *N_i_*, *p_ij_* and *ρ*, where *ρ* is a correlation parameter that accounts for heterogeneity in detection probability. In practice, this solution amounts to using a Binomial distribution with a random *p*, which comes from a Beta distribution. In short, our model combines the biological variation of abundance among roosts with the sampling process of parrot detection:

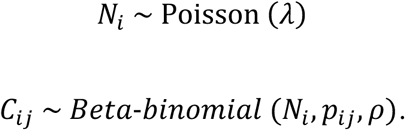

We fit this model to each month’s and to each type of count result (MR or HC) in a Bayesian framework using gamma-distributed vague priors for *λ* and the Beta-binomial parameters (Martin et al. 2011; Supplemental Material Appendix A). The model implementation used the BUGS language (Lunn *et al.* 2000) running on JAGS (Plummer 2003), using code adapted from Kéry and Royle (2015, chap. 6; Supplemental Material Appendix A). To draw from the posterior probability distribution of the parameters, we used an MCMC algorithm with three chains, 25,000 iterations and a burn-in of 5,000 implemented in the software JAGS. All chains converged to R-hat < 1.1.

Even though the detection part of our regional-scale model accounts for variation in detection probability between parrots of different-sized groups, it still assumes constancy of the detection process among sites and counts of the same month. We see this as a reasonable assumption for the regional-scale counts, which were always coordinated by the same individual, consistently applying the same technique. At whole-range scale, however, the sparsity of the data and heterogeneity of counting teams are such that we find it unreasonable to assume constancy of detection parameters. We believe this puts us too close to the limits of applicability of N-mixture models (Barker *et al.* 2017) and we thus present the whole-range count results as raw counts, as they were reported by each team. Because it does note account for unperfect detection, the whole-range count should be interpreted as a lower bound for population size. It is just a lower bound that results from the sum total of counts from all sites, including some of the regional-scale counts from WSC.

In order to explore the implications of imperfect detection on our assessment of total population size, we performed the exercise of extrapolating the estimated density of the Vinaceous-breasted Parrot in WSC to a set of hypothetical geographic ranges of the species. First, we divided the estimate of population size obtained at the regional scale by the WSC area, and then extrapolated the resulting density to three alternative areas that represent a range of possible geographic distributions. The first two areas are the IUCN’s “Extant” and “Possibly Extant” geographic ranges of the Vinaceous-Breasted Parrot (BirdLife International and Handbook of the Birds of the World 2016). The third is the potential range of South American *A. angustifolia* forests as mapped by Hueck (1966) and georeferenced by Hasenack *et al.* (2017). When using population size as listing criteria, the IUCN employs a number of reproductively mature individuals, rather than total population size. Accordingly, we also multiplied the extrapolated population size estimates by an estimated proportion of mature individuals in the total population. Since we are not aware of an empirical assessment of population structure in wild Vinaceous-breasted Parrot populations, we employed two alternative sources for the proportion of mature individuals: the two-thirds undocumented proportion used by the IUCN Red List and an average of the proportions that we could find in the literature for similar-sized, neotropical parrots. The two published sources provide, respectively, a range of 0.17 to 0.34 for *Ara rubrogenys* (Tella *et al.* 2013), and 0.07 to 0.43 for *Amazona autumnalis* (Berg and Angel 2006). They have the same average of 0.25, which we employed as the second proportion of mature individuals.

## Results

The whole-range counts added up to 3,888 and 4,084 individuals, respectively, in 2016 and 2017 (Table 1). Due to logistical constraints, parrots were not counted in 2017 in Argentina. Brazil sites accounted for 93% of individuals in 2016, and 99% in 2017. The total number increased by 5% from the first to the second year, even though there were ten fewer sites visited in 2017 (68 sites) than in 2016 (78). If one accounts only for the sites that were visited in both years (Supplemental Material Table S1), however, the total declines by 15.6%, from 2,938 in 2016, to 2,478, in 2017. The highest number of individuals counted at one site was 356 in 2016 and 364 in 2017. The two counts come from sites approximately 150 km apart, both in the Brazilian state of Santa Catarina and both from the month of May, toward the end of the non-breeding season. Santa Catarina had the highest subtotal count (Table 1), with more than 60% of individuals in both years; followed by Paraná, with approximately 20%, and Rio Grande do Sul, with 8-10%. Santa Catarina was also the state with the highest average number of parrots counted per site, in both years, with 111 individuals per site in 2016 and 174 in 2017.

**Table 1.** Number of Vinaceous-breasted Parrots counted in Argentina, Brazil, and Paraguay during the whole-range counts of 2016 and 2017. Cells with dashes denote changes in the distribution of effort. In 2017 we added point counts in Espírito Santo but were not able to cover Argentina.

| Country | Region | Number Counted |  |
| --- | --- | --- | --- |
|  |  | 2016 | 2017 |
| Argentina | Misiones | 252 | — |
| Brazil | Espírito Santo | — | 2 |
|  | Minas Gerais | 58 | 135 |
|  | Paraná | 803 | 805 |
|  | Rio Grande do Sul | 335 | 409 |
|  | Santa Catarina | 2,324 | 2,606 |
|  | São Paulo | 93 | 109 |
| Paraguay | Alto Paraná | 23 | 18 |
| TOTAL |  | 3,888 | 4,084 |

Comparison of the MR and HC results from counts of the same roost and month reveals that while MR values were always higher, as expected, they were also less variable (Supplemental Material Table S2). Accordingly, when fitting models to MR and HC results separately, estimates of detection probability (*p*) and the precision of abundance estimates (*N*) were generally higher for the MR than for the HC results. We will, for this reason, focus on the MR results in the remainder of the paper. We will refer to MR counts simply as ‘counts’, and specify ‘HC counts’ when we refer to the highly conservative results.

Considering the aggregate of all roosts, we found the lowest number of individuals in the two extremes of the non-reproductive period (Supplemental Material Table S2; Figure 3): in December 2015, with a maximum count of 265 and *N* estimate of 286 ± 8, and in July 2016 with a maximum count of 321 and *N* estimate of 396 ± 22 individuals. The highest aggregate WSC count (1,151 individuals) and *N* estimate (1,393 ± 40 individuals) were obtained in May 2017. In 2016, the maximum aggregate estimate of *N* was 936 ± 40 individuals. The maximum aggregate estimate went up by almost 50% from 2016 to 2017, but if we include only WSC roosts that were counted in both years, the maximum aggregate estimates, with respective standard error and 95% credible interval (c.i.) in square brackets, were 945 ± 52 [859, 1,066] in 2016 and 1,068 ± 44 [994, 1,164] in 2017.

**Figure 3.**
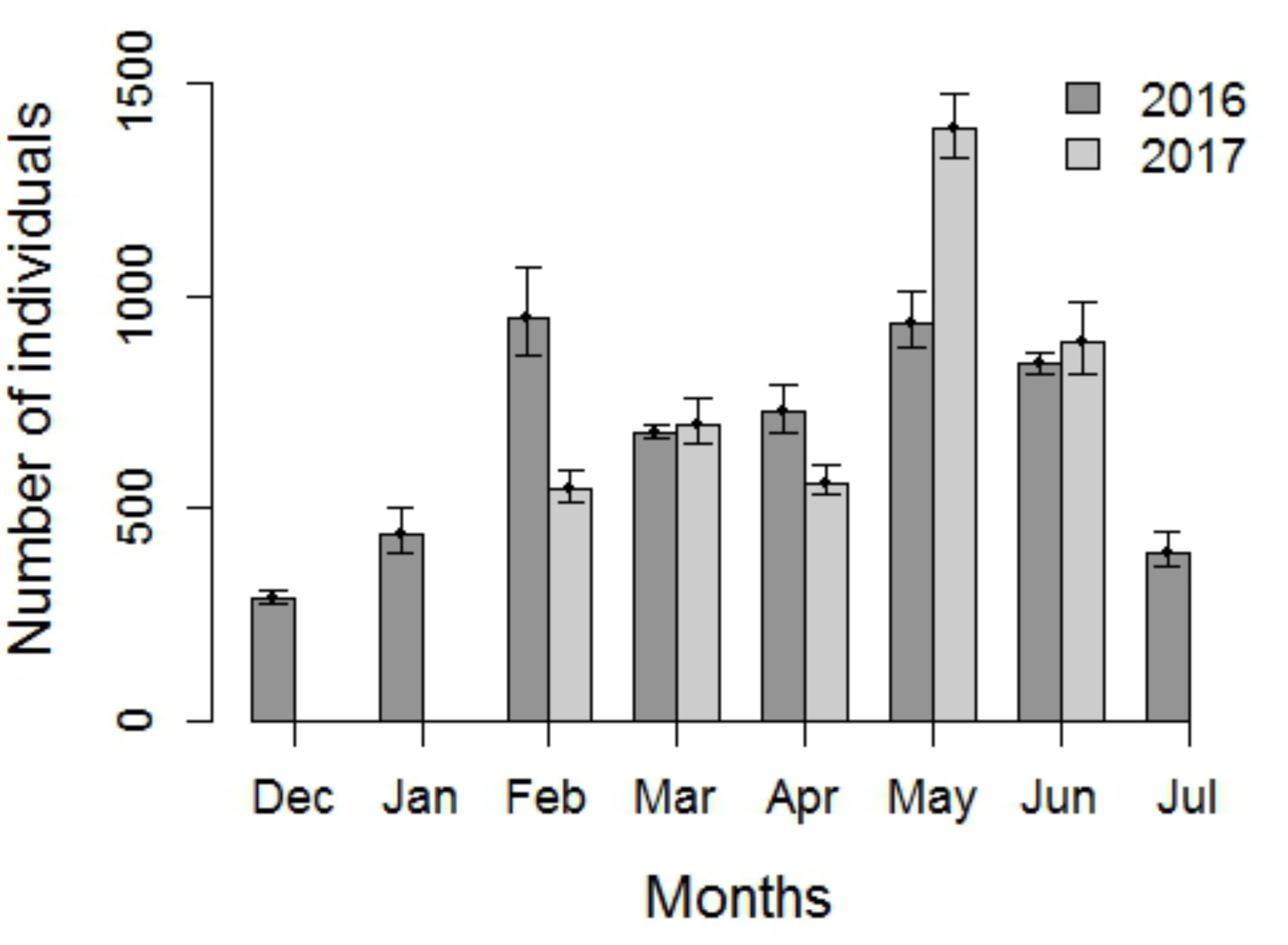
Monthly estimates of the number of Vinaceous-breasted Parrots in WSC in 2016 and 2017 according to the ‘most reasonable’ (MR) count results. Error bars show 95% credible intervals around the estimated number of individuals.

Five of the nine WSC roosts – *Guatambu*, *Ipuaçu*, *Abelardo Luz*, *Água Doce* and *Lebon Régis* - reached *N* estimates in excess of 200 at some point during the sampling period. All roosts showed substantial variation in *N* between months in both years, but there was no obvious synchrony in the temporal variation of the number of individuals at different roosts. The lowest, as the highest *N*, were obtained in different months depending on the roost. For example, while *Água Doce* peaked in March 2016 and May 2017, *Guatambu* did so in April 2016 and February 2017. *Abelardo Luz* was the only roost that peaked both years in the same month, in June. Looking at the spatial distribution of roosts in Figure 2, and the distribution of *N* estimates in Supplemental Material Table S2, it becomes apparent that, in 2016, the northeast of the study area (*Abelardo Luz* and *Água Doce*) concentrated between 56 and 90% of the population during the last three months of the sampling period. However, this tendency was not apparent in 2017, when the same roosts concentrated between 31 and 34% of the population during May and June.

Considering the area of WSC and the maximum aggregate estimate of abundance in each year (*N* = 945 ± 50 [859, 1,066] in February 2016, and *N* = 1,393 ± 40 [1,323, 1,477] in May 2017), we estimate the density of *A. vinacea* in WSC to be between 0.025 and 0.031 individuals per square kilometer in 2016, and between 0.039 and 0.043 in 2017. Extrapolating this density to the areas of the IUCN Extant range (~145,700 km^2^), the IUCN Possibly Extant range (~380,000 km^2^), and to the potential distribution of *Araucaria angustifolia* forests (~175,000 km^2^; Fig. 1), we obtain the values on the first row of Table 3. The numbers for each year are directly proportional to the size of the area, with a minimum of 4,050 individuals for the IUCN Extant range in 2016 (c.i. = [3,681, 4,568]) and a maximum of 15,529 individuals (c.i. = [14,479, 16,466]) for the IUCN Possibly Extant range, in 2017. Multiplying the extrapolated population sizes by the proportions of reproductive individuals obtained from the IUCN Red List and from the scientific literature, we obtained total numbers of reproductive individuals. These values range from 1,012 in 2016, for the IUCN Extant area and a population structure with 25% reproductive individuals, to a maximum of 10,404 in 2017, for the IUCN Possibly Extant area and a population structure with 67% reproductive individuals (Table 3).

**Table 3.** Total number of Vinaceous-breasted Parrots in 2016 and 2017 according to the whole-range counts and to the extrapolation of WSC density for three target areas. The first row shows numbers of individuals of any flying age. The second and third rows show numbers of reproductive individuals according to two different sources of population structure: the IUCN Red List entry for Vinaceous-breasted Parrot, and scientific literature reports of the proportion of reproductive individuals in wild populations of other neotropical parrot species. Values in columns 4-9 of the first row result form multiplying the mean estimate of WSC density by the corresponding extrapolation area. Values in columns 2-9 and rows 2-3 result from multiplying the numbers in row 1 by the corresponding proportion in column 1.

| Proportion of count or extrapolation | Whole-range counts |  | Extrapolations of WSC density for: |  |  |  |  |  |
| --- | --- | --- | --- | --- | --- | --- | --- | --- |
|  |  |  | IUCN Extant range |  | IUCN Possibly Extant range |  | *Araucaria angustifolia* forest |  |
|  | 2016 | 2017 | 2016 | 2017 | 2016 | 2017 | 2016 | 2017 |
| 1.00 | 3,888 | 4,084 | 4,050 | 5,969 | 10,535 | 15,529 | 4,862 | 7,167 |
| 0.67* | 2,605 | 2,724 | 2,713 | 3,999 | 7,057 | 10,404 | 3,257 | 4,802 |
| 0.25† | 972 | 1,016 | 1,012 | 1,492 | 2,634 | 3,882 | 1,216 | 1,792 |

\* Proportion of reproductive individuals in the population of Vinaceous-breasted parrots according to the IUCN Red List (BirdLife International 2017).
† Average proportion of reproductive individuals in wild populations of *Ara rubrogenys* (Tella et al. 2013) and *Amazona autumnalis* (Berg and Angel 2006).

## Discussion

Using two approaches—an extensive range-wide count and an extrapolation from more intensive work at the regional level—we provide evidence that the global Vinaceous-breasted Parrot population consists of a few thousand, but definitely not more than twenty thousand individuals. Considering the extrapolations in Table 3, the 2016 total number of individuals for the IUCN Extant range is the most similar to the whole-range count. This extrapolation exceeds the corresponding whole-range count by 162 individuals, yet its 95% credible interval of 3,681 to 4,568 includes the count by a wide margin. In 2017, the same extrapolation returns 5,969 individuals, well in excess of the whole-range count, which falls outside the extrapolation’s credible interval. Extrapolations for the IUCN Possibly Extant range and for the *A. angustifolia* area are higher than for the IUCN Extant area, in both years. The comparison between approaches reveals information about the various sources of uncertainty inherent to counting parrots at roosts.

The WSC population estimates and the whole-range counts, as well as the extrapolated whole-range estimates, increased from 2016 to 2017, but there is no evidence that this change resulted from population growth. Instead, the increase was largely explained by the addition of two new roosts to the WSC sample, one of them (*Ipuaçu*) with more than 300 individuals estimated for 2017. The WSC estimate increased by almost 50% between years, with non-overlapping credible intervals; however, when we compare yearly estimates based on data from the same seven roosts that were sampled in both years the posterior mean increases by less than 15%, with widely overlapping credible intervals. Likewise, the whole range count, which increased by 4.6% when all sites were summed, decreased by more than 16% when we included only sites that were counted in both years. We conclude that the increase in estimates from 2016 to 2017 is due mostly to improved coverage of the species range and we stress the importance of further improvement.

The IUCN criterion C for the assignment of species to Red List categories states that a species should be considered Endangered if its population is ‘estimated to number fewer than 2,500 mature individuals’ *and* is in decline (BirdLife International 2001). Given the short temporal scope of our study, we do not examine the decline condition, but we can ask whether the 2016 and 2017 population is below the threshold of 2,500 mature individuals. If we multiply the IUCN’s two-thirds (~0.67) proportion of mature individuals by the whole-range results and by the extrapolated numbers of both years, the resulting numbers of mature individuals are always above the IUCN threshold (even though the threshold falls within the credible interval for the 2016 IUCN Extant range value.) The result is very different, though, if we employ the one quarter proportion of reproductive individuals reported by Berg and Angel (2006) and by Tella et al. (2013). In this latter case, four out of six extrapolated population sizes are below the endangerment threshold, with only the values corresponding to the ‘Possibly Extant’ area falling above 2,500 individuals.

These extrapolated population sizes aim to reflect uncertainty about range size, population structure, and local WSC population density. Nonetheless, we caution that these are an insufficient basis for proposing a category change for the species. For one, the numbers are based on the assumption of homogenous density throughout the range, and on a highly uncertain range of population structures. Furthermore, the IUCN assigns threat levels based on a combination of five criteria (Mace *et al.* 2008). In order to qualify for one level, a species must meet conditions from any of the five criteria for that level. Non-fulfillment of the conditions under criterion C would require examination of range and population dynamic conditions under the other criteria, which are beyond the scope of this study. We suggest that the species should remain in the ‘Endangered’ IUCN threat category pending demographic studies and analysis of the conditions under criteria A, B, D and E. Ideally, given appropriate coverage of the species range and understanding of population dynamics, one should be able to assess an extinction risk for the species, which is demanded by criterion E.

Clearly, the assessment of extinction risk can only be as good as the underlying estimates of population size. Our estimate for WSC and its extrapolation are far from perfect, but they show some of the ways in which researchers can address sources of uncertainty in monitoring efforts of Vinaceous-breasted and other parrots. At the broadest level, there is uncertainty about species’ ranges. For Vinaceous-breasted Parrot, we tried to reduce this uncertainty by searching for new roosts 8 days/year in WSC, which returned a 125% increase in the number of sampling sites over the 2 years of the study. We covered the northern half of WSC in more detail than the southern half, which has only one known roost (*Guatambu*; Figure 2). We expected more roosts in the north, because it has more Araucaria forest and a higher density of large (≥ 5 km^2^) forest patches; yet, judging from the distribution of sightings in WikiAves (Wikiaves 2008) and verbal reports, we believe there are more regular roosting sites to be found in the southern part of WSC. Many Vinaceous-breasted Parrots detected in the whole-range counts were also outside the IUCN range, showing that range uncertainty extends well beyond the limits of WSC. The small areas suggestive of isolated populations in the IUCN Extant range (e.g., Figure 1) may be part of larger areas of continuous use, and may be useful starting points for improving knowledge about the species’ distributions.

A second source of uncertainty is the possible variation in density (individuals per unit area) across a species’ range. In our extrapolation of Vinaceous-breasted Parrot abundance from WSC to the whole range, we assumed homogenous density, which is questionable for two reasons. First, densities of organisms tend to be low at the edge of distribution ranges (Brown *et al.* 1995, Gaston 2009). Such a pattern is supported by the relatively lower counts of Vinaceous-breasted Parrots found in Argentina, Paraguay and Espírito Santo (Cockle *et al.* 2007, Segovia and Cockle 2012) when compared with those of eastern Santa Catarina (Prestes *et al.* 2014), which could result in our extrapolation overestimating the true global population size. Secondly, roost density of Vinaceous-breasted Parrots (average number of known roosts per unit area) appears to be lower in WSC (2.6×10^-4^ roosts/km^2^) than over the entire IUCN Extant range (4.0×10^-4^ roosts/km^2^). Thus, if the number of individuals per roost is sufficiently stable across the range, our extrapolation could also be an underestimate of the true global population. Since the whole-range counts do not correct for imperfect detection and estimates of detection probability range from 0.64 to 0.89 (Table 2), the sum of counts is likely to be an underestimate as well. Lacking more robust information about population density outside WSC, we find it reasonable to draw a first estimate of global population size based on the assumption of homogenous density. It is important, however, that this first estimate is taken as what it is—an approximation. Researchers could account for geographic variation in parrot density and improve knowledge of global population size by replicating counts within short periods over several parts of a species’ range. Homogenous or non-homogenous, the spatial distribution of parrots is bound to be dynamic. Such dynamism is unequivocally supported for Vinaceous-breasted Parrots by the disappearance of individuals from roosts during the breeding season and by the variation in WSC roost estimates throughout the study (Supplemental Material Table S2). This brings up a third source of uncertainty, about movement of individuals between roosts, which we tried to address in the present study. We estimated the lowest numbers of Vinaceous-breasted Parrots at all WSC roosts in December 2015 and July 2016 (Table 2)—first and last months of the sampling period of 2016—but the variation of abundance through time was far from synchronous across roosts (Supplemental Material Table S2). Indeed, the estimates for *São Domingos* and *Abelardo Luz* were lowest in January and March of 2016, respectively, both not at the extremes of the sampling period. If there were a gradual accumulation of individuals at all roosts with a peak in the middle of the non-breeding period, we would be inclined to believe that each roost aggregates individuals that breed in the surrounding area. The irregularity of temporal variation in roost size, however, suggests that Vinaceous-breasted Parrots probably move well beyond the immediate surroundings of one roost as they track resources during the non-breeding season (see also Forshaw 2010, Prestes *et al.* 2014). As a result, individuals counted at one roost in a given month, may very well be present at a different roost in another month. This is why we based our WSC estimate on the month with the highest estimate of each year (February 2016 and May 2017) and not on a sum of each roost’s highest monthly estimate. Uncertainty about movement is also the reason behind concentrating counts in as short a period as possible, both in our monthly WSC counts and in the whole-range counts.

**Table 2.** Estimated number of Vinaceous-breasted Parrots (*N*) and their detection probability (*p*), by month, for the aggregate of all roosts sampled in Western Santa Catarina. The numbers in parentheses show aggregate count, based on the sum of the highest count of each roost for the corresponding month. The two rows per month separate estimates based on the ‘most reasonable’ (MR) and the ‘highly conservative’ (HC) count results. Boldface numbers identify the highest *N* estimate of each year.

| Month | 2016 |  | 2017 |  |
| --- | --- | --- | --- | --- |
| | $N$ | $p$ | $N$ | $p$ |
| December (MR) | 286±8 (265) | 0.87±0.03 |  |  |
| (HC) | 275±11 (244) | 0.79±0.03 |  |  |
| January (MR) | 440±28 (335) | 0.69±0.04 |  |  |
| (HC) | 387±28 (297) | 0.67±0.05 |  |  |
| February (MR) | **945±52 (696)** | 0.67±0.03 | 544±19 (426) | 0.69±0.02 |
| (HC) | 954±56 (670) | 0.62±0.04 | 463±18 (374) | 0.71±0.03 |
| March (MR) | 678±9 (639) | 0.89±0.01 | 696±27 (587) | 0.79±0.03 |
| (HC) | 611±7 (588) | 0.92±0.01 | 719±41 (529) | 0.64±0.04 |
| April (MR) | 729±28 (562) | 0.64±0.02 | 559±18 (493) | 0.81±0.03 |
| (HC) | 750±35 (538) | 0.58±0.03 | 498±22 (418) | 0.75±0.04 |
| May (MR) | 936±33 (790) | 0.77±0.03 | **1,393±40 (1,151)** | 0.75±0.03 |
| (HC) | 1006±50 (758) | 0.64±0.03 | 1,195±27 (1,060) | 0.82±0.02 |
| June (MR) | 840±13 (761) | 0.81±0.01 | 891±42 (639) | 0.65±0.03 |
| (HC) | 797±13 (724) | 0.81±0.01 | 917±57 (588) | 0.55±0.03 |
| July (MR) | 396±22 (321) | 0.74±0.04 |  |  |
| (HC) | 353±19 (286) | 0.75±0.04 |  |  |

So far, we discussed three sources of uncertainty that are mostly biological in nature – uncertainty about range limits, about spatial distribution of abundance, and about movements between roosts. Another two sources of uncertainty—double counting (false positive) and imperfect detection (false negative)—are more methodological in nature, but should also guide decisions of study design and data analysis for estimating population sizes in parrots and other birds. In parrot roost counts, double counting happens when observers overestimate the number of parrots in a flock, and when parrots move out of sight and are mistakenly counted as different individuals when they reappear. Our consideration of MR and HC results was an attempt to evaluate the consequences of being less or more conservative about the possibility of double counting. The consequences were negligible: the 95% credible intervals of the MR and HC-based estimates for WSC overlapped in all but six months (March, June and July of 2016; February, April and May of 2017). In those months, the difference was on average 82 individuals, with a standard deviation of 58.6. The tendency for higher precision in MR than HC estimates stems from a greater agreement among MR, than among HC results of the same roost and month. This is no proof that MR counts are indeed closer to the true value, but it does support our reliance on the MR estimates. By including MR and HC estimates in monitoring efforts for other parrots, researchers can assess the potential effects of double-counting on population estimates.

The second methodological source of uncertainty is the recurrent failure to detect some of the parrots that are present at a site. Despite all our efforts to surround the roosts, work with three-observer teams, and connect observers within each team by radio, the counts taken by different teams at the same place and time still differed. We believe that this problem cannot be eradicated, but it should be accounted for. Detection probability (*p*) was always estimated to be greater than 0.6, which is reassuring; however, its variation through time makes it clear that detection probability can’t be estimated once and subsequently used to correct all counts from then on. Researchers can address the fact of *p* < 1 by replicating counts and estimating *p* during every time period for which they want to estimate *N*. It should be noted that even though part of the field team gained experience with the species, the sites, and the logistics over the course of the study in WSC, *p* did not increase monotonically from the beginning to the end of the sampling period. Instead, *p* varied from month to month without any apparent trend, reaching its maximum in March 2016 and its minimum in April 2016 (Table 2). This suggests that failure to detect parrots at roost counts is not just a matter of observer experience, but largely a matter of chance, weather, and unpredictable parrot movements.

While considering the variation of detection probability and its effects on estimates of population size, it is important to take a critical look at our choice of modeling roost count data with an N-mixture approach. N-mixture models are a useful tool for obtaining estimates of abundance from counts of unmarked animals (Royle 2004). Barker et al. (2017) argue that heterogeneity in detection probability among counts creates issues of parameter identifiability in N-mixture models, seriously limiting the modeler’s ability to estimate abundance. Kéry (2018) investigates these issues over three different statistical distributions of abundance among sites, finding virtually no evidence of identifiability problems under a Poisson distribution. Faced with a group of species and a spatial scale of work for which mark-recapture would be unfeasible, we still find the N-mixture approach the most appropriate option for analyzing our data. We do emphasize, however, the importance of taking sampling design precautions for limiting heterogeneity of detection probability among counts. When opting among statistical distributions to model variation in abundance among sites, the choice is not trivial but so far the Poisson seems to offer the best predictions, even if not always the best fit (Kéry 2018).

Habitat loss and nest poaching have caused obvious but poorly documented declines of many Neotropical parrot populations, including Vinaceous-breasted Parrots (Wright *et al.* 2001, Ribeiro *et al.* 2009, Berkunsky *et al.* 2017). Any efforts to protect these species will benefit from improved knowledge of population size and structure. We hope that our approach to estimating population size of Vinaceous-breasted Parrots in WSC and beyond will motivate others to obtain replicated counts of parrot roosts for this and other species. In an attempt to coordinate observers and gather count information for Vinaceous-breasted Parrots, we set up an online count-reporting tool where users can access existing data and contribute their own. The current version is available in Portuguese at: http://vivianezulian.azurewebsites.net/. The uncertainty surrounding regional- and whole-range population estimates, however, is still high enough to justify monitoring Vinaceous-breasted Parrots, and other Neotropical parrots, with a wide variety of observation techniques. On one front, citizen science networks such as WikiAves, Xeno-Canto, and eBird can offer valuable information for mapping species ranges and reproductive areas. On the other, molecular analysis of parrots across their range would help understand seasonal movements and the spatial structure of populations. Progress on both fronts will require formal integration of different types of data into one statistical model of species distribution and abundance. Any progress on the molecular front will require employment of effective and safe techniques for obtaining parrot DNA without endangering the sampled individuals. Our study illustrates several key sources of uncertainty about parrot abundance estimates, and how they can be addressed through monitoring protocols and statistical analysis. Critically, by addressing and estimating uncertainty, parrot monitoring efforts can move beyond minimum or average roost counts to a broader understanding of what we do and do not know about parrot numbers, eventually producing reliable assessments of population trends over time.

## Acknowledgments

We are deeply grateful to the 112 observers and local coordinators who counted *Amazona vinacea* throughout their distribution in 2016 and 2017. We thank Juarez Camara and Deizi Groth for invaluable local assistance and encouragement in WSC. *FATMA*, the *Grimpeiro* nongovernmental organization, *Palmasola* S/A lumber company, *Ministerio de Ecología y RNR* (Misiones), and many local landowners facilitated access to field sites or housing. We also thank IUCN for species range shapefiles.

Funding statement: Fieldwork was supported by Aves Argentinas/AOP, the Loro Parque Foundation, and by grants from the Rufford Foundation (18013-D to KLC and 19835-1 to VZ), *Fundação Grupo Boticário de Proteção à Natureza* (to JM), Columbus Zoo & Aquarium (to KLC), and CNPq (PP 312606/2013-3, to GF).

Ethics statement: We followed the Ornithological Council’s Guidelines to the Use of Wild Birds in Research.

Author contributions: GF, VZ, ESM and KLC conceived the idea of the study. VZ and ESM supervised roost counts in WSC, KLC in Argentina, AL in Paraguay. JM, NPL, and RTJ coordinated the whole-range counts. All authors collected field data. GF supervised the WSC research and quantitative analysis carried out by VZ. GF, VZ, and KLC wrote the paper.

